# The rifampicin-inactivating mono-ADP-ribosyl transferase of *Mycobacterium smegmatis* significantly influences reactive oxygen species levels in the actively growing cells

**DOI:** 10.1101/2020.01.10.902668

**Authors:** Sharmada Swaminath, Atul Pradhan, Rashmi Ravindran Nair, Parthasarathi Ajitkumar

## Abstract

A classic example of antibiotic inactivating function in bacteria is the *Mycobacterium smegmatis* (*Msm*) encoded rifampicin-inactivating mono-ADP-ribosyl transferase (*arr*). Since its probable biological role has been proposed to be in DNA damage response, which is inflicted by reactive oxygen species (ROS), in the present study, we examined whether *Msm* Arr influences ROS levels. For this purpose, the levels of the ROS, hydroxyl radical and superoxide, were determined in the mid-log phase (MLP) cells of *Msm arr* knockout (*arr*-KO) strain, in comparison to those in the equivalently grown *Msm arr*^+^ wild-type (WT) strain. The MLP *arr*-KO cells generated significantly elevated levels of superoxide and hydroxyl radical, unlike the equivalently grown WT MLP cells. Complementation of *arr*-KO with *arr*, but not with empty vector, restored the ROS levels comparable to those in the WT strain. Elevated ROS levels in the *arr*-KO strain enabled selection of rifampicin-resistant mutants at 10^-7^ cfu/ml from the rifampicin-unexposed MLP cells of *arr*-KO, which is one-log_10_ higher than that for WT cells (10^-8^). Upon prolonged exposure to rifampicin, the susceptibility, persister formation, generation of elevated levels of hydroxyl radical by the persisters, rifampicin-resister generation frequency of the persisters and regrowth of the rifampicin-resistant mutants from the respective persisters were all comparable between the *arr*-KO and WT strains. These observations revealed that Arr influences ROS levels in the actively growing *M. smegmatis* cells but not in the rifampicin-exposed cells. We proposed the probable pathway through which Arr might be influencing ROS levels in the actively growing *M. smegmatis* cells.

**IMPORTANCE:** Diverse genera of bacteria consisting of pathogens, opportunistic pathogens and non-pathogens, possess Arr-type activities that confer equally efficient rifampicin resistance, thereby posing serious health hazard. Acquisition of this function by other bacteria through horizontal gene transfer enhances the hazard posed by the bacteria possessing it. *M. smegmatis* is an opportunistic human pathogen that causes infections of skin and soft tissues. Moreover, *M. smegmatis* is a genetically tractable model organism for *M. tuberculosis* with the potential to function even as tuberculosis vaccine. In view of these significant aspects of Arr and *M. smegmatis*, the study to find out the natural physiological role of Arr in *M. smegmatis*, gains importance for designing strategies to prevent antibiotic inactivation and to target the cellular function to contain the bacterium. Above all, the three-dimensional structure of *M. smegmatis* Arr reveals significant structural homology with eukaryotic ADP-ribosyltransferases and bacterial toxins, thereby giving the study broad significance.

## Introduction

Bacteria of diverse genera, constituting pathogens, opportunistic pathogens and non-pathogens, use covalent modification of antibiotics as one of the many strategies to inactivate antibiotics (reviewed in 1-6). These modifications include ribosylation, phosphorylation, acylation, glycosylation, nucleotidylation, and thiol transfer (reviewed in 2, 6). The latest addition to the large list of different kinds of modification is the inactivation of a broad range of rifamycins by the class A flavoprotein monooxygenase coded for by a drug-inducible *rox* gene of *Streptomyces venezuele* (*rox-sv*) (7). It monooxygenates the naphthyl group of rifamycin causing permanent inactivation of the antibiotic (7). Rifamycin has been found to get modified by ADP-ribosylation also, conferring rifamycin resistane (8). The presence of rifamycin-inactivating function in the soil bacterium *S. venezuele* is of great benefit in destroying the wide range of rifamycins polluting the environment. However, the presence of antibiotic-inactivating functions in environmental bacteria can also be a potential hazard, considering the emergence of antibiotic-resistant bacterial strains by horizontal gene transfer across bacterial genera (reviewed in 9). In fact, comparison of the genetic organisation of the *arr* locus of several bacterial genera, including that of *M. smegmatis* (*Msm*), suggested historic acquisition of the gene through horizontal gene transfer (8).

Mono-ADP-ribosylation activity, being one of the types of covalent modifications of antibiotics and coded for by mono-ADP-ribosyltransferases, holds the double purpose of antibiotic inactivation and a cellular role in bacterial physiology (reviewed in 10, 11). ADP-ribosylation has been found to be involved in several cellular functions in many organisms from bacteria to humans (reviewed in 12). The *arr* genes have been found in various genera of pathogenic and non-pathogenic bacteria, conferring equally efficient rifampicin-resistance (13–15). The product of the *arr* gene, MSMSEG_1221, mono-ADP-ribosylates rifampicin thereby rendering it inactive (16–18). This function is the major contributor to rifampicin tolerance by *Msm*. Further, the rescue of rifampicin-mediated transcription inhibition by MsRbpA (19) is a minor contributor to rifampicin tolerance.

In addition to ADP-ribosylation of rifampicin, *Msm* Arr is known to ADP-ribosylate two endogenous proteins of 30 kDa and 80 kDa, the identities of which have not yet been established (20). However, the Arr in *Streptomyces coelicolor* A3(2), the SCO2860 protein, homologous to *Msm* Arr (reviewed in 21), ADP-ribosylates four endogenous proteins, BldKB (67 kDa), MalE (45 kDa), a putative periplasmic branched-chain amino acid-binding protein (42 kDa), and a putative periplasmic solute binding protein (40 kDa) (22). In *Msm*, an extracellular binding protein (489 residues, ∼55 kDa), a sugar-binding protein (∼45 kDa), and a D-xylose binding periplasmic protein (∼40 kDa) have been found to be the homologues of the BldKB (67 kDa), MalE (45 kDa), and the periplasmic solute binding protein (40 kDa), respectively, based on BLAST alignment (23). Due to their homologies, it was suggested that these proteins in *Msm* could be the substrates of *Msm* Arr, with the biochemical confirmation pending (23).

Arr-2, a protein sharing 55% identity with *Msm* Arr, which is present in several transposons and integrons of Gram-negative pathogenic bacteria (24–27), is believed to confer resistance to rifampicin. The *Msm* Arr does not possess sequence homology to known protein ADP-ribosyl transferases such as mono-ADP-ribosyl transferase (ART), poly ADP-ribosyl transferases (PARPs), or bacterial toxins (8). Despite the lack of amino acid sequence homology to the Arr proteins of diverse bacteria and eukaryotic counterparts, the *Msm* Arr shares significant three-dimensional structural homology with important ADP-ribosyltransferases in eukaryotes, including poly(ADP-ribose) polymerases (PARPs) and bacterial toxins (8). This common feature among these proteins gives broad significance to the study of the cellular function of *Msm* Arr.

Mono-ADP-ribosylation has been found to modify different macromolecules in the cells, which in turn affects many key biological processes in all the organisms, from bacteria to humans (reviewed in 10, 21). These cellular processes include transcription, DNA replication, DNA-damage repair, signal transduction, cell division, response to infection, stress response, microbial pathogenicity, and aging (21). In *Msm*, the substrate binding and catalytic activities of *Msm* Arr were found to be required for the regulation of a small subset of DNA damage responsive genes, in addition to conferring rifampicin resistance (28). Further, the catalytic-activity-impaired *Msm* Arr was also required for the suppression of a set of ribosomal proteins and rRNA during DNA damage and stress response (28). The transcription of *Msm arr* gene was found to be responsive to oxidative stress and DNA damage, which suggested that Arr is involved in general stress response (28).

*Msm*, isolated from environment and human smegma, is an opportunistic pathogen that causes specific types of infections in blood, skin, soft tissues and/or bone in humans (29–33). Many of the isolates from humans are resistant to isoniazid and rifampin but susceptible to ethambutol, doxycycline, sulfamethoxazole, ciprofloxacin, imipenem, and amikacin (29). Besides the pathological importance of *Msm*, it is considered to be a genetically tractable model organism for *Mycobacterium tuberculosis*, with the proposed potential of a genetically modified *Msm* as a vaccine against *M. tuberculosis* (34). Meanwhile, the study of the antibiotic-inactivating function is critical to design strategies to make the antibiotic effective against the bacterium, study of its physiological role is important to devise approaches to eliminate the opportunistic bacterium *per se*.

## Results

### The experimental design and strategy

*Msm* Arr was proposed to be required for DNA damage response (28), which is known to be triggered by ROS inflicted DNA damage (reviewed in 35). Therefore, the objective of the present study was to find out whether the natural physiological role of *Msm* Arr in the actively growing mid-log phase (MLP) populations of the cells involves influence of ROS levels and thereby resister generation frequency inflicted by them in the bacterium against rifampicin in the antibiotic-unexposed condition and upon prolonged exposure to lethal concentrations of the antibiotic. In order to achieve this objective, an *arr* deletion knockout mutant (*arr*-KO) was first generated. The growth characteristics of the *arr*-KO strain were then compared to those of the *arr+* wild type (WT) strain to find out whether there were any gross growth differences between them. Subsequently, the minimum bactericidal concentration (MBC) of rifampicin was determined for the *arr*-KO strain. Then, the levels of the reactive oxygen species (ROS), hydroxyl radical and superoxide, were determined in the mid-log phase (MLP) cells of *arr*-KO, in comparison to those in the equivalently grown WT, and in the *arr*-complemented *arr*-KO and the empty-vector-complemented *arr*-KO strains, which were constructed as single copy genome integrants.

The levels of specific reactive oxygen species (ROS), hydroxyl radical and superoxide, were first examined in the actively growing mid-log phase (MLP) cells of these four strains, stained with dyes specific for the two ROS, followed by determination of the levels of fluorescence of the oxidised dyes using flow cytometry. The levels of hydroxyl radical were determined in the WT and *arr*-KO strains using electron paramagnetic resonance (EPR) spectrometry also. The frequency of emergence of resisters selectable with rifampicin from the actively growing MLP cultures of *arr*-KO and WT strains was determined by plating rifampicin-unexposed cultures on rifampicin plates. Further, the response of the *arr*-KO and WT strains to rifampicin upon prolonged exposure was examined, in terms of their extent of susceptibility, rifampicin persister formation, generation of elevated levels of hydroxyl radical by the persisters, frequency of the generation of rifampicin-resisters from the respective persisters and regrowth of the mutation-gained persisters. The nature of these responses of the *arr*-KO strain were compared to those of the WT strain and the naturally *arr*-lacking *M. tuberculosis* exposed to rifampicin for prolonged duration, which was earlier reported by us (36). Such comparison of *arr*-KO with WT revealed that the natural physiological role of *arr* lies in influencing ROS levels in the actively growing cultures of *Msm*. The probable pathway of its influence on ROS levels was then predicted.

### Generation and growth characteristics of the *arr*-KO mutant

The mono-ADP-ribosyl transferase (*arr*; MSMEG_1221) knockout mutant (*arr*-KO) was generated using the allelic exchange method, as described (37), with minor modifications. The recombination event was confirmed using polymerase chain reaction (PCR) using two primer sets, one set with the primer specific for the gene upstream of *arr* and reverse primer for hygromycin resistance (*hyg^r^*) gene (selection marker) and the other set with the forward primer of *hyg^r^* and the reverse primer specific for the gene downstream of *arr* (Fig. 1A, B). The stability of the *arr*-KO mutant was verified and confirmed by culturing it in the absence of hygromycin and subsequently in the presence of hygromycin for 12 generations each. Growth curve experiments showed that the WT and *arr*-KO strains have comparable mass doubling time of 2.8 ± 0.22 hrs and 3.1 ± 0.11 hrs, respectively (Fig. 1 C-E). Thus, the deletion of the *arr* gene did not affect the overall growth of the strain. This stable *arr*-KO mutant strain, which grows like the WT strain, was used in all the experiments.

**FIG 1.**
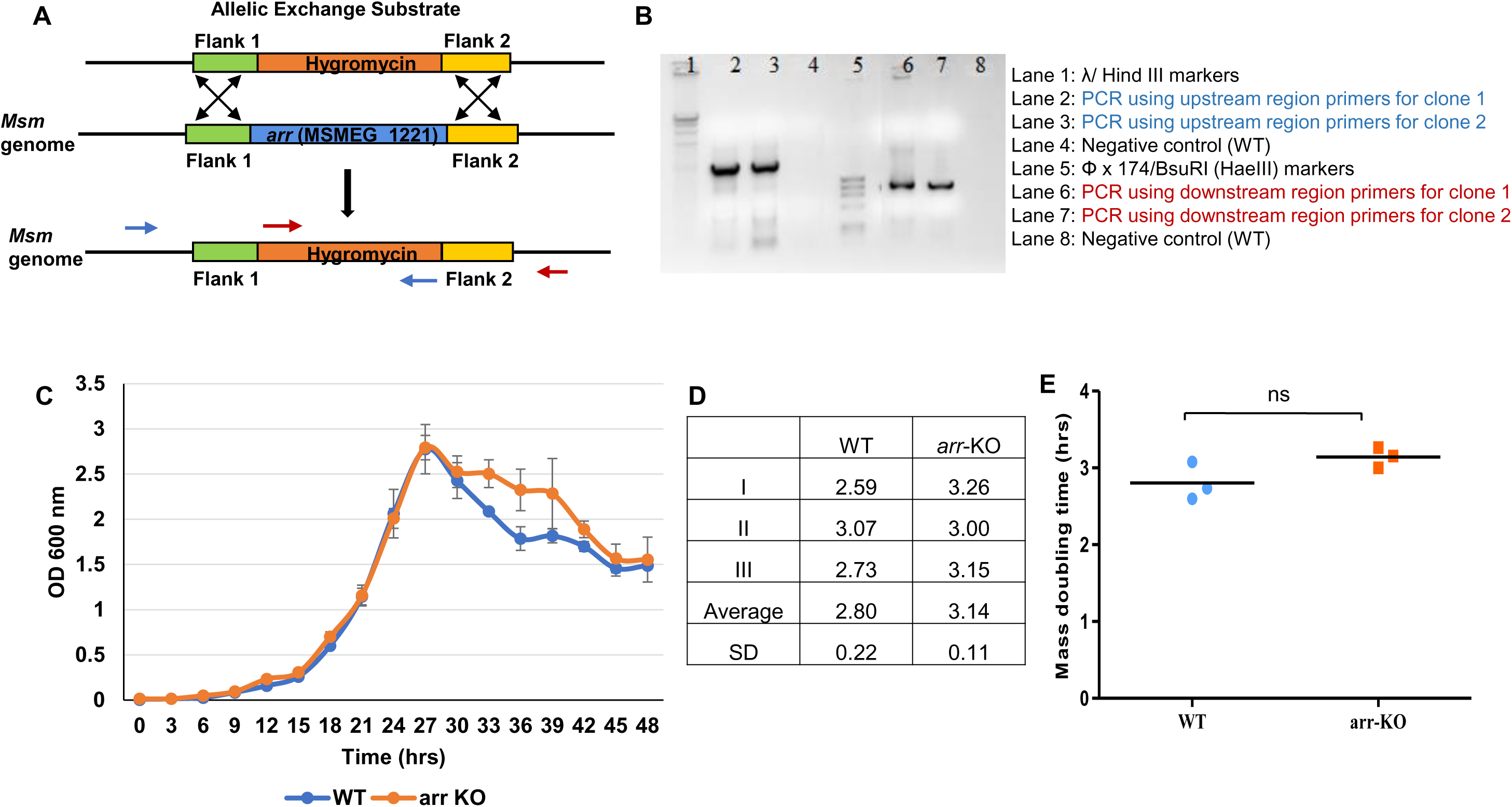
Generation of *Msm arr*-KO strain and comparison of growth characterstics of *Msm* WT and *arr*-KO strains. (A) Diagrammatic representation of the homologous recombination of allelic exchange substrate used for the generation of the *arr*-KO strain. (B) Confirmation of the recombination event by PCR using primers upstream (blue primers in Fig. 1A) and downstream (red primers in Fig. 1A) of the recombination locus. (C) Growth curves and (D) mass doubling time (calculated from growth curves) of WT and *arr*-KO strains from the respective biological triplicates. (E) Comparison of the mass doubling time of the strains. The statistical significance was calculated using two-tailed unpaired t-test. ns, indicates no significance.

### Significantly elevated levels of hydroxyl radical in the MLP cells of *arr*-KO

Several studies have shown that both the ROS, hydroxyl radical and superoxide, contribute to oxidative stress and consequential mutagenesis (DNA damage) in bacterial cells (reviewed in 35; 38, 39). Since *Msm* Arr has been implicated in DNA damage reponse (28), we first examined the MLP cells of *arr*-KO and WT for the presence of elevated levels of hydroxyl radical, using two different approaches: (i) electron paramagnetic resonance (EPR) spectrometry, the gold standard method for the measurement of hydroxyl radical (40, 41), and (ii) flow cytometry measurement of the fluorescence emission of hydroxyphenyl fluorescein (HPF), a hydroxyl radical specific dye (42), by the respective HPF-stained cells.

#### Using EPR spectrometry

Analysis of the lysates of the MLP cells of *arr*-KO and WT using EPR showed the characteristic strong signal specific to 5,5-dimethyl-1-pyrroline N-oxide (DMPO)-OH adduct from the lysates of the *arr*-KO cells, unlike the weak signals from the lysates of the WT cells (Fig. 2A, B). The intensity of the signal specific to DMPO-OH adduct in the lysates of *arr*-KO cells was significantly higher by ∼2-fold in comparison to that of WT cells (Fig. 2C). It indicated that the absence of *arr* generated high levels of hydroxyl radical.

**FIG 2.**
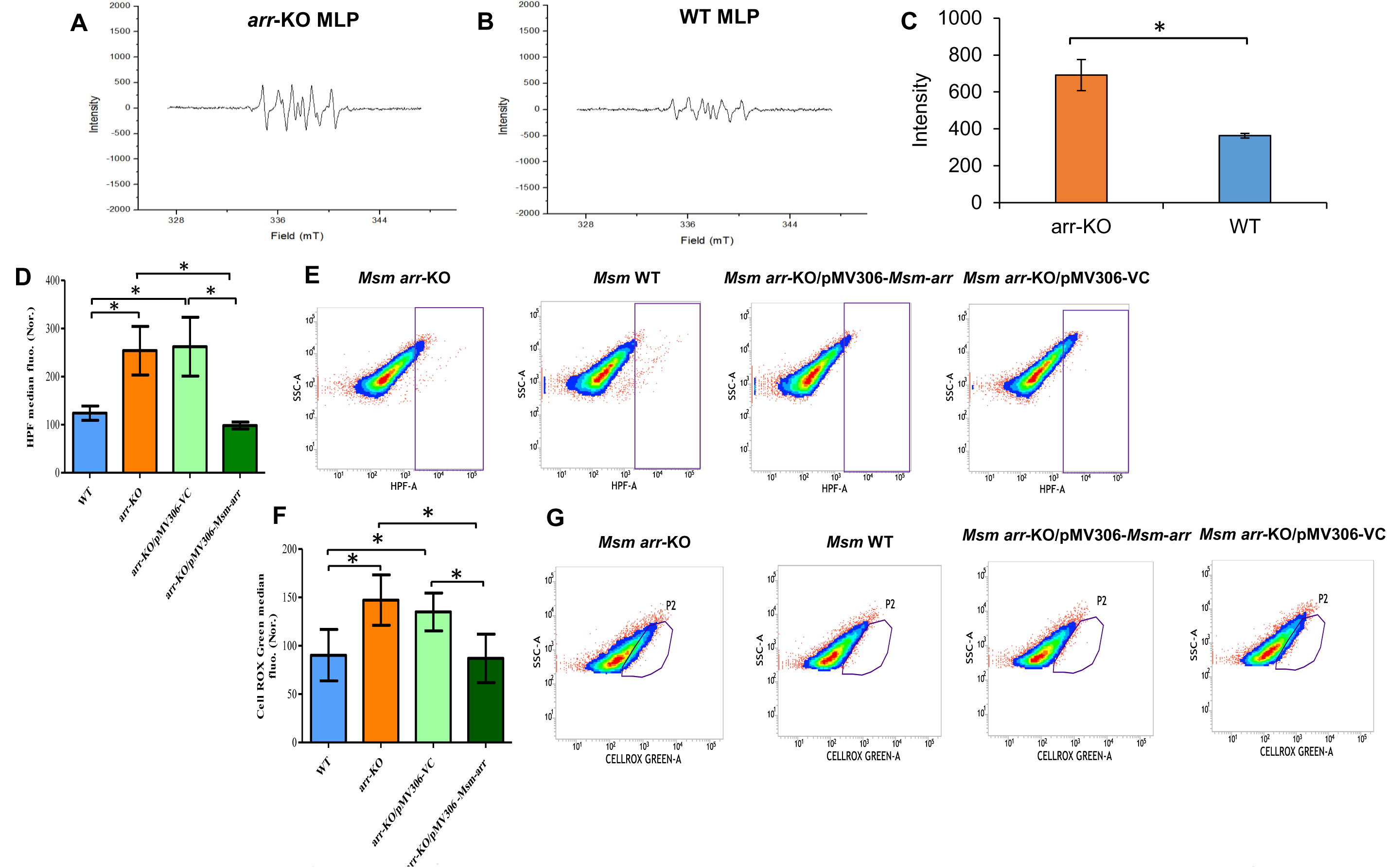
Detection of hydroxyl radical and superoxide in the MLP cells of *Msm* WT and *arr*-KO using electron paramagnetic resonance spectrometry (EPR) ytometry. (A-B) Representative EPR spectra of DMPO-OH adduct from lysates of (A) *Msm arr*-KO and (B) *Msm* WT MLP cells. (C) Quantitative MPO-OH adduct in the lysates of the *Msm arr*-KO MLP cells and *Msm* WT MLP cells (n = 3). (D-G) Flow cytometry profiles of *Msm* WT and *arr*-O/pMV306-*Msm*-*arr* and *arr*-KO/pMV306-VC strains, stained with HPF for hydroxyl radical and CellROX Green for superoxide. (D) The average an fluorescence normalised with its respective autofluorescence for the MLP cells of the strains (n = 3). (E) Representative density plots of HPF ce from the MLP cells of the strains. (F) The average CellROX Green median fluorescence normalised with its respective autofluorescence for ells of the strains (n = 3). (G) Representative density plots of CellROX Green fluorescence from MLP cells of the strains. In (C, D, F) one asterisk s P value less than or equal to 0.05 (P ≤ 0.05). The statistical significance was calculated using two-tailed paired t-test.

#### Using flow cytometry

For the flow cytometry analysis, an *arr*-complemented *arr*-KO strain and empty-vector-complemented *arr*-KO strain (control), which were single copy genome integrants, were also constructed for comparison (Fig. S1). In agreement with the EPR data, the HPF-stained *arr*-KO MLP cells showed significantly elevated levels of fluorescence as compared to those in the WT MLP cells (Fig. 2D, E). Complementation of the *arr*-KO strain with the genome-integrated pMV306-*Msm-arr*, which carries *arr* gene under its own promoter (Fig. S1), significantly reduced the HPF fluorescence to a level comparable to that of the WT cells, indicating the restoration of hydroxyl radical levels comparable to those in the WT cells (Fig. 2D, E). Complementation with the vector control plasmid (pMV306-VC) did not reduce the HPF fluorescence of the *arr*-KO cells (Fig. 2D, E). Thus, the elevated levels of hydroxyl radical in the absence of *arr* and its restoration to normal levels upon complementation with *arr*, but not with the empty vector, indicated the influence of hydroxyl radical levels by *arr* in *Msm* cells.

### High levels of superoxide in the MLP cells of *arr*-KO than in the WT

It is known that hydroxyl radical is generated by Fenton reaction between hydrogen peroxide and labile Fe^2+^ ions (43). Further, majority of hydrogen peroxide is formed by the dismutation of superoxide by superoxide dismutase (44). The Fe^2+^ ions for Fenton reaction is obtained by the leaching of labile iron from 4Fe-4S proteins by superoxide (45). Therefore, the presence of significantly elevated levels of hydroxyl radical in the *arr*-KO MLP cells implied the possibility of the presence of elevated levels of superoxide in them. This possibility was verified by an assay using the superoxide-specific dye, CellROX Green (46, 47), by staining the MLP cells of WT, *arr*-KO, *arr*-KO/pMV306-*Msm*-*arr* (*arr* complemented *arr*-KO), and *arr*-KO/pMV306-VC (vector control) strains and measuring the fluorescence of the oxidised dye. For preventing any reduction of superoxide by redox-active metals, diethylenetriaminepentaacetic acid (DTPA) was added to inactivate redox-active metal ions (48, 49). The MLP cells of the *arr*-KO and *arr*-KO/pMV306-VC strains showed significantly high levels of fluorescence indicating significantly high levels of superoxide (Fig. 2F, G). On the contrary, the MLP cells of the WT and the *arr*-complemented *arr*-KO/pMV306-*Msm*-*arr* showed significantly lower and comparable levels of fluorescence, indicating lower levels of superoxide as compared to the levels in the *arr*-KO strain (Fig. 2F, G). These observations confirmed the presence of elevated levels of superoxide in the MLP cells of the *arr*-KO strain, indicating that the superoxide levels might be influenced by *arr* in *Msm* cells.

### MBC of rifampicin for the MLP cells of *arr*-KO and WT

High levels of superoxide and hydroxyl radical are known to cause DNA damage by inflicting mutations (reviewed in 35; 38). Since the *arr*-KO cells produced high levels of ROS, the rifampicin-resister generation frequencies of the actively growing *arr*-KO MLP cells unexposed to rifampicin and upon prolonged exposure to rifampicin were determined. For this purpose, the 1x MBC of rifampicin for 10^8^ cells/ml of *arr*-KO was first determined to be 2.08 µg/ml. This was significantly (∼20-fold) lower than the 1x MBC of rifampicin (42 µg/ml) for the WT reported recently by us (50; Fig. S2A and B). However, the 1x MBC of moxifloxacin, against which *arr* does not confer tolerance (antibiotic control), was determined to be 0.125 µg/ml for *arr*-KO (Fig. S2C) and 0.133 µg/ml for WT, as reported (50), which were comparable (Fig. S2D). Thus, the MBCs of moxifloxacin for the WT and *arr*-KO strains were not different, and the reduction was found only in the MBC of rifampicin for *arr*-KO, as compared to that for WT. This confirmed the specificity of the activity of Arr, which is known to inactivate rifampicin only (15, 17), thereby specifically influencing the MBC of rifampicin only, and not that of moxifloxacin.

### Rifampicin-resister generation frequency of the MLP cells of *arr*-KO and WT

We determined the rifampicin-resister generation frequency of the MLP cells of the *arr*-KO and WT strains against the respective 3x MBC of rifampicin by directly plating the MLP cells, cultured unexposed to rifampicin, on 3x MBC rifampicin plates. The *arr*-KO MLP cells showed variation in the resister generation frequency, as compared to those of WT, against the respective 3x MBC of rifampicin (n = 9 for each sample) (Fig. 3A, B). Although the resister generation frequency of *arr*-KO was most often higher than that of WT, there was no significant difference between the values of *arr*-KO and WT (Fig. 3A, B). However, when they were exposed to their respective 4x and 5x MBCs of rifampicin, the resister generation frequency of the WT strain decreased progressively and significantly, but not that of the *arr*-KO strain (Fig. 3C). Interestingly, the resister generation frequency of the *arr*-KO MLP cells continued to remain comparably higher but not significantly different against the 3x, 4x and 5x MBCs of rifampicin (Fig. 3C). The lack of change in the resister generation frequency of the *arr*-KO MLP cells against 3x, 4x, and 5x MBC of rifampicin hinted that the absence of Arr in the *arr*-KO strain seemed to have caused loss of some sort of control on the response of the cells to increasing concentrations of MBC of rifampicin. This observation correlated with the significantly elevated levels of superoxide and hydroxyl radical in the actively growing rifampicin-unexposed MLP cells of the *arr*-KO strain but not in the equivalent cells of the WT strain (see Fig. 2).

**FIG 3.**
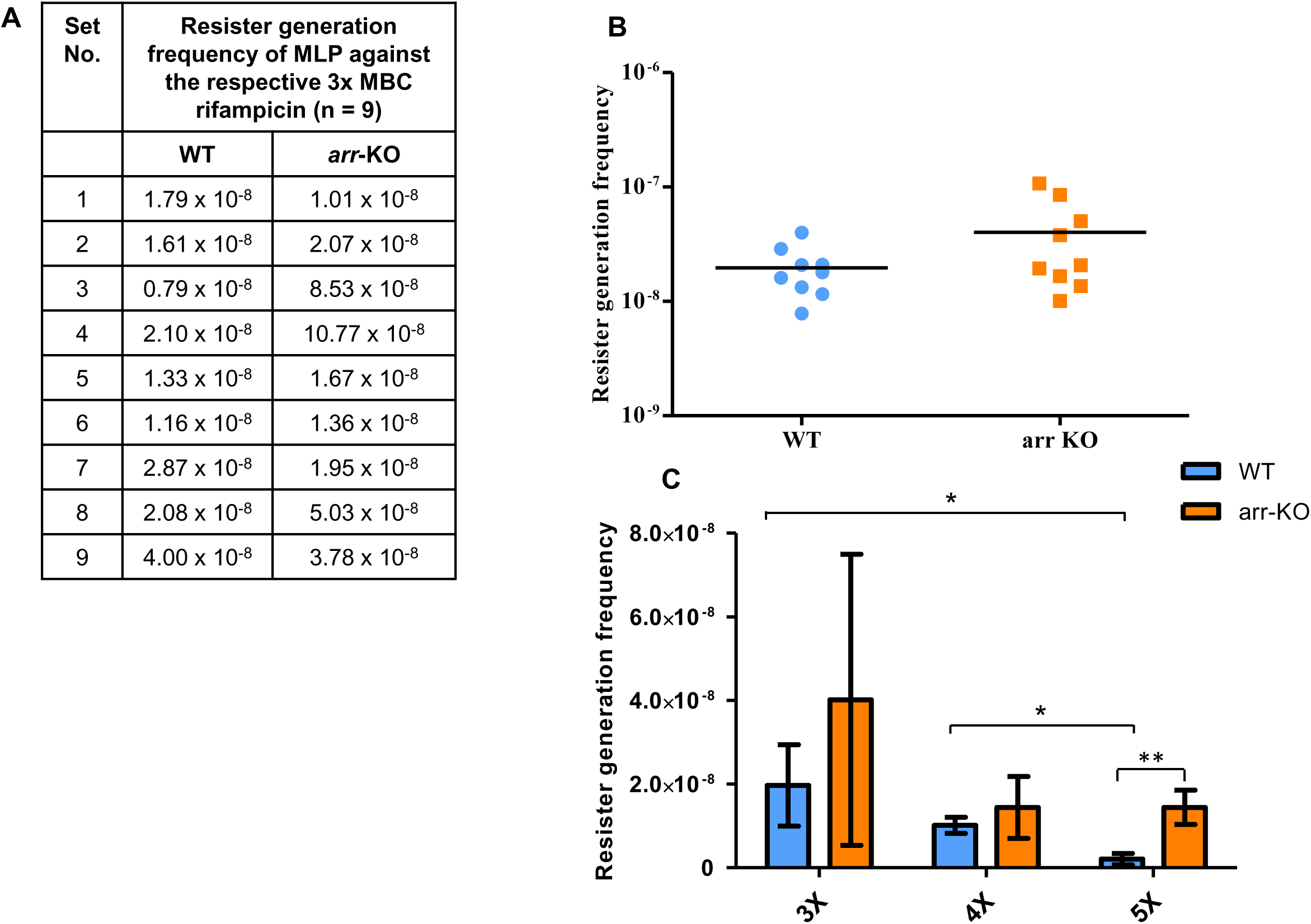
Rifampicin-resister generation frequency of the MLP cells of *Msm* WT and *arr*-KO strains. (A, B) Resister generation frequency against respective 3x MBC of rifampicin. (A) Table of values and (B) scatter plot of resister generation frequency for MLP cells of *Msm* WT and *arr*-KO strains against the respective 3x MBC rifampicin (n = 9). (C) Comparative rifampicin-resister generation frequency of the MLP cells of *Msm* WT and *arr*-KO strains against the respective 3x (plotted from the values in Fig. 3A), 4x and 5x MBC of rifampicin (n = 9, for 3x and n = 3 for 4x and 5x MBC). In Fig (C), one asterisk (*) indicates P value less than or equal to 0.05 (P ≤ 0.05) and two asterisks (**) indicate P value less than or equal to 0.01 (P ≤ 0.01). Statistical significance was calculated using two-tailed unpaired t-test and with Welch’s correction for comparing unequal sample sizes.

### The response of *arr*-KO and WT to rifampicin upon prolonged exposure

We had recently demonstrated that prolonged exposure of *M. tuberculosis* MLP cells to MBC of rifampicin and moxifloxcin and of *Msm* to moxifloxacin caused the formation of the respective antibiotic persisters (36, 50). These antibiotic persisters generated elevated levels of hydroxyl radical, underwent genome-wide mutagenesis and emerged *de novo* at very high frequency as the respective antibiotic-resistant genetic mutants. From these persisters specific to the antibiotics, genetic mutants that were resistant to other antibiotics could also be selected, which was an outcome of the genome-wide mutagenesis by the elevated levels of ROS (36, 50). In view of these observations, the *arr*-KO and WT strains were exposed to high MBC of rifampicin for prolonged duration to find out whether the absence of *arr* would cause any alteration in the response of the *arr*-KO mutant to rifampicin upon proloned exposure or behave in a manner similar to the WT strain.

#### (i) Killing, persistence, and regrowth of rifampicin-exposed WT and *arr*-KO

The MLP cells of WT and *arr*-KO were exposed to the respective ∼2x MBC of rifampicin for 96 hrs in biological triplicate cultures of cell density 10^8^ cells/ml. The CFU/ml of the WT and *arr*-KO strains were determined at regular intervals by plating aliquots of the respective cultures on rifampicin-free plates. The response of the WT and *arr*-KO cells to their respective MBC of rifampicin involved sequential killing, persister, and regrowth phases, as in the case of the response of *M. tuberculosis* to MBC of rifampicin and moxifloxcin and of *Msm* to moxifloxacin (Fig. 4A, red line; 4B, brown line, respectively; 36, 50). The killing phases of the WT and *arr*-KO cells were characterised by a steady decrease of about 3-log_10_ unit and 4-log_10_ unit of CFU/ml, respectively, on rifampicin-free plates (Fig. 4A, red line; 4B, brown line, respectively). The killing phase was ensued by the persister phase, where no appreciable change was observed in the CFU/ml of the WT and *arr*-KO (Fig. 4A, red line; 4B, brown line, respectively). The persistence phase was followed by the regrowth phase showing a notable increase in the CFU/ml on rifampicin-free plates (Fig. 4A, red line; 4B, brown line, respectively).

**FIG 4.**
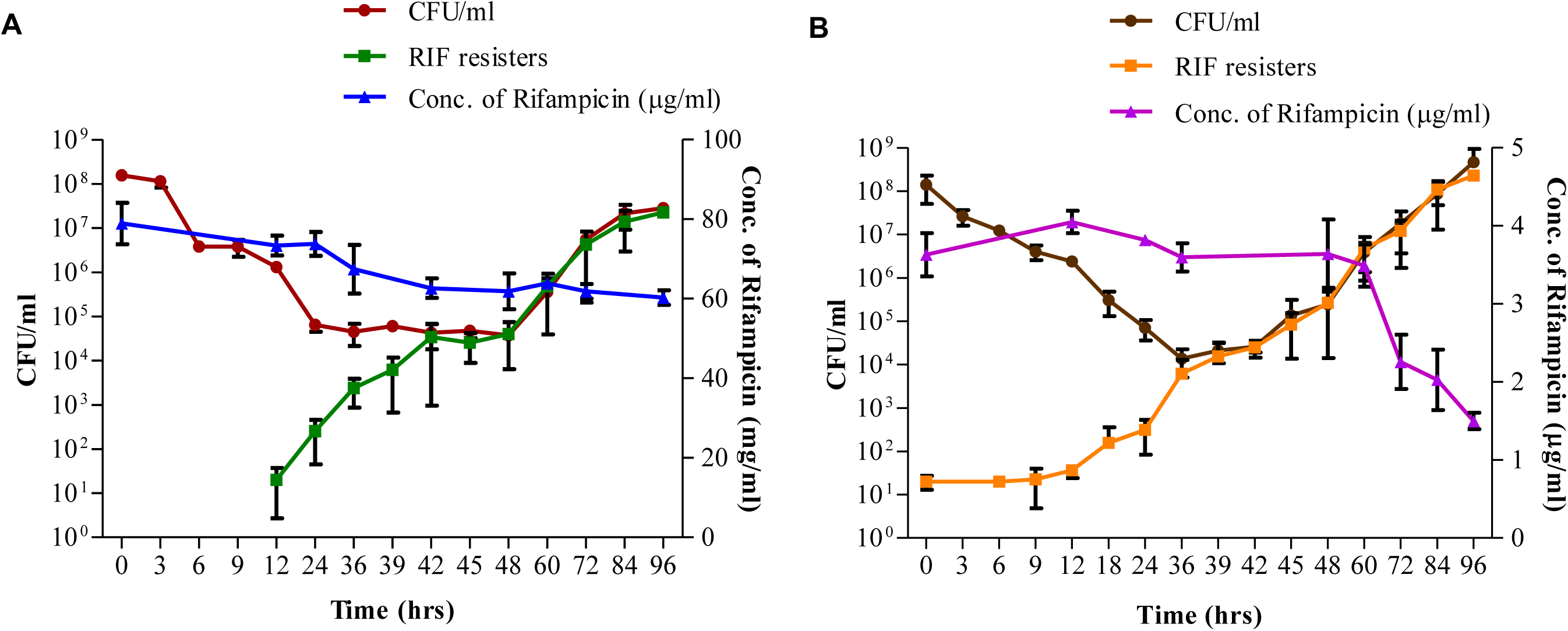
Rifampicin-susceptibility profile of the MLP cells of the *Msm* WT and the *arr-*KO strains upon prolonged exposure to rifampicin and the emergence of rifampicin-resisters during the exposure. (A) Rifampicin-susceptibility profile of the WT strain exposed to ∼2x MBC (75 µg/ml) of rifampicin for 96 hrs and plated on rifampicin-free plate (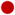, red line), the CFU/ml on ∼3x MBC (125 µg/ml) rifampicin plates (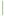, green line), and the concentration of rifampicin during the course of the experiment (right *y* axis; 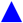, blue line) (n = 3). (B) Rifampicin-susceptibility profile of the *arr-*KO strain exposed to ∼2x MBC (4 µg/ml) of rifampicin for 96 hrs and plated on rifampicin-free plates (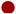, brown line), the CFU/ml on ∼3x MBC (6 µg/ml) rifampicin plates (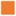, orange line) and the concentration of rifampicin during the course of the experiment (right *y* axis; 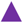, purple line) (n = 3).

The rifampicin concentration in the Middlebrook 7H9 medium, as determined using rifampicin bioassay (51), showed only a slight decrease from the original concentration in the respective cultures (Fig. 4A, blue line; 4B, purple line, respectively). However, in the case of *arr*-KO, a steep reduction in the rifampicin concentration was observed from the 60 hr of exposure, with ∼1.5 µg/ml (0.75x MBC) of rifampicin still present even at 96 hr of exposure (Fig. 4B, purple line). The regrowth of the WT and *arr*-KO strains, as found from the CFU/ml on the rifampicin-free plates, obtained by plating the aliquotes of the cultures growing in the continued presence of lethal concentrations of rifampicin indicated that these cells might be rifampicin-resistant *rpoB* mutants (52-54, reviewed in 55). Similar regrowing population was found by us in the case of rifampicin/moxifloxacin-exposed *M. tuberculosis* cells and moxifloxacin-exposed *Msm* cells for prolonged duration (36, 50). The regowing cells might also be rifampicin-tolerant phenotypic variants surviving by many other multiple mechanisms known (56, 57, reviewed in 58).

#### (ii) Emergence of rifampicin-resisters from the regrowth phase

The CFU/ml obtained from plating aliquots of the rifampicin-exposed cultures of WT and *arr*-KO from 0 hr onwards on rifampicin plates, containing 125 µg/ml and 6 µg/ml rifampicin (∼3x MBC equivalent rifampicin for WT and *arr*-KO, respectively), confirmed the emergence of rifampicin-resistant mutants from the respective regrowth phase cultures (Fig. 4A, green line; 4B orange line, respectively). The number of rifampicin-resisters steadily increased during the persister phase (24 hr to 48 hr of exposure to rifampicin in the WT strain and 36 to 42 hr of rifampicin exposure in the *arr*-KO strain) and continued to increase further during the regrowth phase till 96 hr monitored (Fig. 4A, green line; 4B orange line, respectively). The overlapping nature of the CFU/ml in the rifampicin-containing and rifampicin-free plates, from 48 hr and 42 hr, for the WT and the *arr*-KO strains, respectively, indicated that probably all the cells that were regrowing in the continued presence of MBC of rifampicin (∼60 µg/ml; 1.5x MBC in the WT culture and ∼3.5 µg/ml; 1.5x MBC in the *arr*-KO culture) might essentially be rifampicin-resistant genetic mutants carrying mutations in the *rpoB* locus (52-54, reviewed in 55) or rifampicin-tolerant phenotypic variants surviving by multiple mechanisms known (56, 57, reviewed in 58).

#### (iii) Emergence of rifampicin-resisters from the unexposed arr-KO cells

Surprisingly, in the case of the *arr*-KO strain, the rifampicin-resisters could be selected from the 0 hr of exposure itself, at a frequency of 10^-7^ CFU/ml (Fig. 4B, orange line). The CFU/ml of the rifampicin-resisters from the rifampicin plates of *arr*-KO remained more or less the same during the first 9 hrs of exposure and subsequently started increasing dramatically and overlapped with the CFU/ml on the rifampicin-free plates from about 36 hr of exposure onwards (Fig. 4B, overlapping orange line and brown line from 36 hrs). The selection of rifampicin-resisters from the 0 hr (MLP) of rifampicin exposure of the *arr*-KO culture, at a frequency of 10^-7^ CFU/ml, indicated the natural mutation rate of *arr*-KO resulting in the selection of the mutants resistant to rifampicin (Fig. 4B, orange line). Since elevated levels of ROS is one of the reasons for the generation of genetic mutants, the presence of rifampicin-resistant mutants, even prior to the exposure of the *arr*-KO strain to rifampicin, was reflective of the elevated levels of hydroxyl radical in the MLP cells of the *arr*-KO strain (see Figs. 2A-C, D).

On the contrary, in the case of WT, the rifampicin-resisters started emerging only by about 12 hr of exposure, prior to the persister phase (which began at 24 hr of exposure) at a frequency of <10^-7^ CFU/ml (Fig. 4A, green line). The natural rifampicin-resister frequency of *Msm* WT cells was reported to be 10^-8^ CFU/ml (59). Thus, the striking difference that could be noted between the WT and *arr*-KO strains during prolonged exposure to rifampicin was the propensity of the *arr*-KO cells for the generation of rifampicin-resistant natural mutants (compare Fig. 4A, green line with orange line in 4B), at approximately one-log_10_-fold higher frequency than that of WT. This higher level of rifampicin resister generation frequency from the 0 hr of exposure itself was probably reflective of the inherently elevated levels of ROS (superoxide and hydroxyl radical) in the MLP *arr*-KO cells, unlike in the case of MLP WT cells (see Fig. 2).

#### (iv) *arr*-KO rifampicin resisters from the rifampicin-unexposed MLP cells and rifampicin persister cells

The formation of rifampicin-resistant colonies on high MBC (∼3x MBC) rifampicin plates (see Fig. 4A, green line; 4B, orange line) indicated the possibility of the presence of genetic mutations conferring rifampicin resistance. The rifampicin resistance of 95% of the rifampicin-resistant *M. tuberculosis* strains has been found to be due to mutations in the **R**ifampicin **R**esistance **D**etermining **R**egion (**RRDR**) of *rpoB* (Fig. S3; 52-54, reviewed in 55). Sequencing of the RRDR of the *rpoB* from the 12 rifampicin resisters of the *arr*-KO strain, which were isolated from the 0 hr of rifampicin exposure, and from the 30 hr, 36 hr and 39 hr of the persister phase, showed oxidative stress specific C→T or A→G mutations (60, 61, reviewed in 35), resulting in His442-Arg, Arg444-Cys and Ser438-Leu changes (Table 1). The 12 rifampicin-resistant mutants isolated from the 12 hr (killing phase), 24 hr, 30 hr, 39 hr, and 48 hr (from persister phase) of the WT strain also showed C→T or A→G mutations, resulting in His442-Tyr and His442-Arg changes (Table 2). Identical RRDR mutations at identical positions have been observed in the rifampicin-resistant clinical strains of *M. tuberculosis* (52-54, reviewed in 55) and in the *M. tuberculosis* cells regrowing from the rifampicin persister population *in vitro* (36). Calculation of the rifampicin-resister generation frequencies of the persister cells of WT and *arr*-KO strains from the CFU/ml values from the rifampicin plates and rifampicin-free plates in the Fig. 4A and B, respectively, showed high resister generation frequency of the order of 0.394 ± 0.171 and 0.578 ± 0.2581, respectively. These rifampicin-resister generation frequencies of the WT and *arr*-KO persister cells were comparable, but were ∼8-log_10_-fold higher than the resister generation frequencies of their respective MLP cells, which was strongly indicative of very high levels of hydroxyl radical in these persister cells.

**TABLE 1.** List mutations at the RRDR*^a^* of *rpoB* of rifampicin-resisters of *Msm arr*-KO

| Resister<sup>b</sup><br>Name | Mutation<br>(codon) | Nucleotide<br>position | Amino acid<br>change | Aminoacid<br>position<br>in *Msm*)<br>(in *Mtb*)<sup>c</sup> | Type of<br>mutation |
| --- | --- | --- | --- | --- | --- |
| KO-R1-0-1 | CGT-TGT | 1330 | R-C | 444 (447) | C-T |
| KO-R2-0-1 | CGT-TGT | 1330 | R-C | 444 (447) | C-T |
| KO-R3-0-1 | CGT-TGT | 1330 | R-C | 444 (447) | C-T |
| KO-R3-0-2 | CAC-CGC | 1325 | H-R | 442 (445) | A-G |
| KO-R3-0-3 | CAC-CGC | 1325 | H-R | 442 (445) | A-G |
| KO-R1-30-1 | CAC-CGC | 1325 | H-R | 442 (445) | A-G |
| KO-R1-36-1 | CAC-CGC | 1325 | H-R | 442 (445) | A-G |
| KO-R1-39-1 | CAC-CGC | 1325 | H-R | 442 (445) | A-G |
| KO-R2-36-1 | CAC-CGC | 1325 | H-R | 442 (445) | A-G |
| KO-R2-39-1 | CAC-CGC | 1325 | H-R | 442 (445) | A-G |
| KO-R3-36-1 | CAC-CGC | 1325 | H-R | 442 (445) | A-G |
| KO-R3-36-2 | TCG-TTG | 1313 | S-L | 438 (441) | C-T |

<sup>a</sup>RRDR, Rifampicin Resistance Determining Region.
<sup>b</sup> Resisters were named in the order of the culture amongst the triplicates, the hour from which they were isolated and the colony number.
<sup>c</sup>Amino acid position in *Mtb* is given in parenthesis. Note: For reporting the amino acid position change in *Mtb* references *E. coli* numbering is used and 441, 445, 447 amino acid position of *Mtb* corresponds to 522, 526, 528 of *E. coli*, respectively.

**TABLE 2.** List of the mutations at the RRDR*^a^* of *rpoB* of rifampicin resisters of *Msm* WT

| Resister<sup>b</sup><br>Name | Mutation<br>(codon) | Nucleotide<br>position | Amino acid<br>change | Aminoacid<br>position<br>in *Msm*)<br>(in *Mtb*)<sup>c</sup> | Type of<br>mutation |
| --- | --- | --- | --- | --- | --- |
| R1-12-1 | CAC-TAC | 1324 | H-Y | 442 (445) | C-T |
| R2-12-1 | CAC-TAC | 1324 | H-Y | 442 (445) | C-T |
| R2-12-2 | CAC-TAC | 1324 | H-Y | 442 (445) | C-T |
| R2-24-1 | CAC-CGC | 1325 | H-R | 442 (445) | A-G |
| R2-24-2 | CAC-CGC | 1325 | H-R | 442 (445) | A-G |
| R2-24-3 | CAC-CGC | 1325 | H-R | 442 (445) | A-G |
| R2-24-4 | CAC-CGC | 1325 | H-R | 442 (445) | A-G |
| R1-30-2 | CAC-TAC | 1324 | H-Y | 442 (445) | C-T |
| R2-39-1 | CAC-TAC | 1324 | H-Y | 442 (445) | C-T |
| R2-39-2 | CAC-TAC | 1324 | H-Y | 442 (445) | C-T |
| R2-39-3 | CAC-TAC | 1324 | H-Y | 442 (445) | C-T |
| R3-48-3 | CAC-TAC | 1324 | H-Y | 442 (445) | C-T |

<sup>a</sup>RRDR, Rifampicin Resistance Determining Region.
<sup>b</sup>Resisters were named in the order of the culture amongst the triplicates, the hour from which they were isolated and the colony number.
<sup>c</sup>Amino acid position in *Mtb* is given in parenthesis. Note: For reporting the amino acid position change in *Mtb* references *E. coli* numbering is used and 445 amino acid position of *Mtb* corresponds to 526 of *E. coli*.

### Elevated levels of hydroxyl radical in the rifampicin persisters of *arr*-KO & WT

The oxidative stress specific nature of the mutations, C→T or A→G, in the rifampicin-resistant mutants of the respective persister cells of WT and *arr*-KO indicated that they might have been brought about by the high levels of superoxide and hydroxyl radical. Superoxide is known to facilitate mutagenesis in an indirect manner, when it gets dismutated to H_2_O_2_ (through superoxide dismutase) and also cause leaching of labile iron, which together generate hydroxyl radical by Fenton reaction (43, 45, 62). On the contrary, hydroxyl radical is known to induce mutations directly in a sequence non-specific manner (reviewed in 35; 39). Therefore, it was possible that the high rifampicin-resister generation frequency of the persister cells of the rifampicin-exposed WT and *arr*-KO strains might have been due to the presence of elevated levels of hydroxyl radical in them. In fact, we had earlier reported elevated levels of hydroxyl radical in the rifampicin persisters of *M. tuberculosis* inflicting genome-wide mutations (36). In view of this possibility, we examined the rifampicin persister cells of *Msm* WT and *arr*-KO for the presence of hydroxyl radical, using EPR spectrometry and flow cytometry of HPF-stained persister cells.

#### Using EPR spectrometry

Analysis of the lysates of the rifampicin persister cells of the WT and *arr*-KO strains from the 36 hr of rifampicin exposure showed the characteristic strong signal specific to 5,5-dimethyl-1-pyrroline N-oxide (DMPO)-OH adduct indicating the presence of high levels of hydroxyl radical (Fig. S4A, C). The respective strong signal corresponding to hydroxyl radical in the WT and *arr*-KO lysates was quenched by the presence of the hydroxyl radical quencher, thiourea (TU; 63) (Fig. S4B, D). Quantitation of the strength of the EPR signals of the respective persister cells showed significantly higher levels of hydroxyl radical as compared to the levels in the presence of TU, confirming the specificity of detection of hydroxyl radical (Fig. 5A). These observations confirmed the presence of elevated levels of hydroxyl radical in the rifampicin persister cells of WT and *arr*-KO. It was interesting to note that the rifampicin persisters of *arr*-KO produced higher levels of hydroxyl radical than the WT persisters, although the difference was not statistically significant (Fig. 5A). It indicated that Arr might not have influence on the levels of hydroxyl radical generated in response to rifampicin by the persister cells. Thus, the influence of Arr on ROS levels seemed to be confined to the actively growing *Msm* cells unexposed to rifampicin (see Fig. 2A-C).

**FIG 5.**
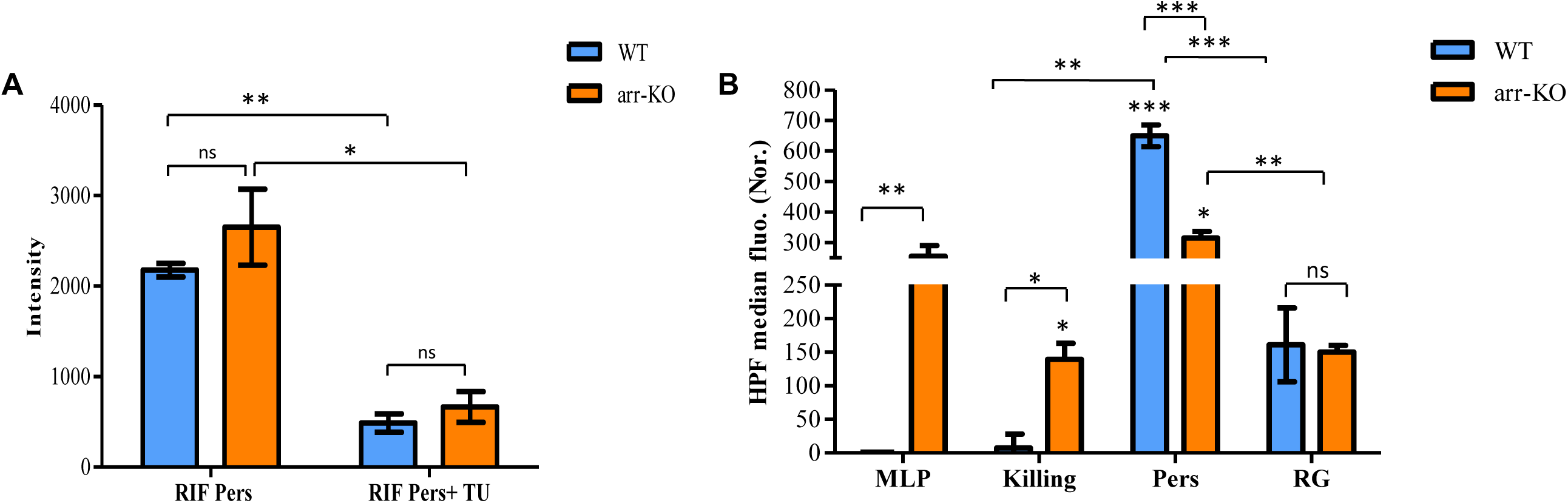
Detection of hydroxyl radical in the *Msm* WT and the *arr*-KO strains exposed to the respective ∼2x MBC of rifampicin using EPR and by flow cytometry. (A) The levels of DMPO-OH adduct in the cell lysates of rifampicin persister cells (RIF Pers), and thiourea-treated rifampicin persister phase (RIF Pers+TU) cells of the WT and *arr-*KO strain, detected using EPR (n = 3, in each case). (B) The average HPF median fluorescence, normalised with its respective autofluorescence, from the MLP, killing, persister (Pers) and regrowth (RG) phases of the rifampicin-exposed cells of the WT and *arr-*KO strain (n = 3, in each case). One asterisk (*) indicates P value less than or equal to 0.05 (P ≤ 0.05), two asterisks (**) indicate P value less than or equal to 0.01 (P ≤ 0.01) and three asterisks (***) indicate P value lesser than 0.001 (P < 0.001). The statistical significance was calculated using two-tailed paired t-test. In (B) the significance denoted on top of the bars are with respect to its corresponding MLP.

#### Using flow cytometry

We first confirmed that the MLP cells of WT and *arr*-KO, which were used for HPF staining, showed killing, persister and regrowth phases upon prolonged exposure to rifampicin (Fig. S5A and C). The flow cytometry profile of the rifampicin-exposed HPF-stained cells of WT and *arr*-KO from the killing, persister, and regrowth phases showed high fluorescence in the respective persister phase cells of both the strains and in the HPF-stained equivalent number of rifampicin-unexposed MLP cells of the *arr*-KO strain (Fig. S5B and D). Quantitation of the flow cytometry data showed that unlike the WT MLP cells, the *arr*-KO MLP cells showed significantly high levels of hydroxyl radical (Fig. 5B), which was reminiscent of the data in Fig. 2C, D. Thus, prolonged exposure of the WT and *arr*-KO strains to the respective MBCs of rifampicin generated persister cells with significantly elevated levels of hydroxyl radical in both the strains. However, the levels of hydroxyl radical in the WT persister cells seemed to be significantly higher than those in the *arr*-KO persister cells (Fig. 5B). This was unlike in the case of the specific detection of hydroxyl radical levels using EPR spectrometry, wherein *arr*-KO cells showed higher levels of hydroxyl radical than the WT cells, although the difference was not significant (compare Fig. 5A and 5B). The higher levels of HPF fluorescence in the WT persister cells in flow cytometry could be probably due to the non-specific binding of HPF to hypochlorite (^-^OCl), as suggested (42), which might be present in the persister cells. Exposure of the persister cells of WT and *arr*-KO to rifampicin beyond the persister phase led to a decrease in the HPF fluorescence, implying low levels of hydroxyl radical, probably due to the emergence of rifampicin-resistant mutants which are no more sensitive to the antibiotic (Fig. 5B; Fig. S5B and D overlays). Thus, the responses of the *arr*-KO and WT strains to the respective MBC of rifampicin upon prolonged exposure were by and large comparable. This observation implied the strong possibility that the natural physiological role of Arr might be confined to the influence of ROS levels in the actively growing *Msm* cells and not in the rifampicin-exposed cells.

## Discussion

### The physiological role of Arr in *M. smegmatis*

In our attempt to find out the natural physiological role of *Msm* Arr, the present study revealed that Arr influences ROS levels in the actively growing cells but not the ROS levels in the rifampicin-exposed cells. It is understandable that its natural physiological role is confined to the cells during active growth, rather than during exposure to rifampicin, as Arr needs to be mostly utilised for the ADP-ribosylation of rifampicin upon exposure to the antibiotic. These conclusions were validated by several observations in the present study: (i). significantly high levels of ROS (superoxide and hydroxyl radical) in the MLP cells of *arr*-KO strain, even without exposure to rifampicin; (ii). restoration of the ROS levels to normalcy upon complementation with *arr* but not with the empty vector; (iii). the 10^-7^ CFU/ml frequency of the emergence of rifampicin-selectable *rpoB* mutants from the rifampicin-unexposed actively growing *arr*-KO MLP cells as compared to the 10^-8^ CFU/ml of the WT strain; (iv). comparability of the nature of the responses of the *arr*-KO and WT strains upon prolonged exposure to MBC of rifampicin, with both the strains showing killing, persiseter and regrowth phases; and (v). comparability of the rifampicin resister generation frequencies of the rifampicin persister cells of the *arr*-KO and WT strains.

The higher levels of hydroxyl radical in the WT strain, as compared to the *arr*-KO strain, in the HPF fluorescence measurement, but the contrary situation in the measurement using EPR spectrometry, can be reconciled with based on the following observations: (i). the higher HPF fluorescence in the WT strain might have been due to the non-specific binding of HPF to hypochlorite (^-^OCl), as suggested (42), which might be present in the persister cells of the WT strain; (ii). EPR spectrometry is more specific for hydroxyl radical determination than the fluorescence of oxidised HPF due to the influence by hypochlorite (^-^OCl); and (iii). the rifampicin resister generation frequencies of the *arr*-KO and WT persister cells, caused by hydroxyl radical, were comparable, despite higher HPF fluorescence in the hydroxyl radical measurement in the WT strain. Thus, these observations indicated that the hydroxyl radical levels of the persister populations of the *arr*-KO and WT strains could be taken to be not significantly different despite the higher HPF fluorescence indicative of hydroxyl radical in the WT strain than in the *arr*-KO strain. Taken together, all the distinct features of the *arr*-KO strain, as compared to those of the WT strain, seemed to suggest that the role of Arr in the influence on ROS levels might be confined to actively growing population and not to the ROS levels in the rifampicin-exposed cells. At present, it is difficult to predict the possible means through which Arr may be influencing ROS levels in the actively growing cells. Nevertheless, it may be speculated that Arr may be mono-ADP-ribosylating critical components in the pathways governing respiratory, redox, or antioxidant systems, and thereby regulating their activities and, in turn, influencing the ROS levels.

### Implications of higher levels of superoxide and hydroxyl radical in *arr*-KO

Superoxide is known to get dismutated to hydrogen peroxide (H_2_O_2_) (44). It is also known to elevate free-iron levels by causing leaching of labile iron from 4Fe-4S proteins (45). The labile iron and H_2_O_2_ undergo Fenton reaction to produce hydroxyl radical (43, 45, 62). Therefore, it was quite possible that the higher levels of hydroxyl radical in the *arr*-KO strain might have been generated by Fenton reaction from the H_2_O_2_ produced from superoxide through dismutation and the labile iron leached from 4Fe-4S proteins by the superoxide, as reported in some Gram-negative bacteria (64, 65). Further, it is known that elevated levels of hydroxyl radical are known to cause genome-wide mutations in bacterial systems, including mycobacteria, from which mutants that are resistant to any antibiotic can be selected (36, 50, 65). This implied that the resister generation frequency of *arr*-KO, which produced significantly higher levels of ROS than the WT strain, would most likely be higher than that of WT. The one-log_10_-fold difference in the rifampicin resister generation frequency between the actively growing MLP cells of *arr*-KO (10^-7^) and WT (10^-8^) was in agreement with this implication. Nevertheless, the high levels of the mutagenic ROS, hydroxyl radical, in the *arr*-KO cells found in our study were consistent with the suggested role for Arr in DNA repair-related events (28), which also involves ROS. Thus, the findings in the present study, on the possible biological role of Arr in influencing ROS levels in the actively growing *Msm*, establish a more clear link with the earlier proposed activity of Arr on DNA damage response (28).

### Striking differences in the response of *Msm arr*-KO strain and the naturally *arr*-lacking *M. tuberculosis* to rifampicin

Even though *M. tuberculosis* naturally lacks *arr*, we noticed several differences between the response of *Msm arr*-KO and *M. tuberculosis* to rifampicin. First, the MBC values of rifampicin for the *Msm arr*-KO strain and *M. tuberculosis* were 2.08 µg/ml and 0.1 µg/ml (36), respectively. Thus, the MBC of rifampicin was 20-fold lower for *M. tuberculosis*, indicating the significantly higher susceptibility of *M. tuberculosis* to rifampicin, unlike the *arr*-KO strain. Hence a similar difference could be expected between their minimum inhibitory concentrations (MICs) as well. Secondly, the *Msm arr*-KO strain inherently generated significantly high levels of hydroxyl radical, unlike *M. tuberculosis*, as reported (36). Thirdly, the sequential phases of killing, persister and regrowth of the cells of the *arr*-KO strain exposed to rifampicin for prolonged duration was similar to the response of *M. tuberculosis* and *Msm* cells exposed to MBC of rifampicin and moxifloxacin, respectively, for prolonged duration (36, 50).

An earlier study using saturated cultures of *Msm arr*-KO strain suggested that the *Msm arr*-KO strain could be used as a model system for anti-tuberculosis drug testing for rifampicin analogues either alone or in combination with other drugs (66).

This suggestion was made based on the comparability of the MIC values of rifampicin for the saturated cultures of *arr*-deletion mutant strain, grown to saturated OD values, and for the naturally *arr*-lacking *M. tuberculosis* (66). However, one may note the conspicuous 20-fold difference in the 1x MBC values of rifampicin for the *Msm* arr-KO (2.08 µg/ml in the present study) and of *M. tuberculosis* (0.1 µg/ml for 10^8^ cells/ml; 36) and the strikingly higher (one-log_10_-fold; 10^-7^ CFU/ml) rifampicin resister generation frequency of the actively growing cultures of the *Msm arr*-KO strain and of *M. tuberculosis* (10^-8^ CFU/ml; 36) to rifampicin. These conspicuous differences between the actively growing *Msm arr*-KO strain and *M. tuberculosis* caution that the actively growing *Msm arr*-KO may not be a surrogate system to test for rifampicin analogues either alone or in combination with other anti-tuberculosis drugs. Further, saturated bacterial cultures suffer from nutritional stress, which contributes to higher oxidative stress and consequential incurrence of mutations, as reported for many bacterial systems, including mycobacteria (67–69).

## Materials and Methods

### Bacterial strains and culture conditions

*Mycobacterium smegmatis* mc^2^155 (*Msm*) (70; Table S1) was used to determine the response of the bacilli to rifampicin. *Msm* was cultured in Middlebrook 7H9 broth (Difco) in containing 0.2% glycerol and 0.05% Tween 80 and grown under shaking condition at 170 rpm in a bacteriological incubator shaker at 37°C until the culture reached **m**id-**l**og **p**hase (**MLP**; OD_600_ nm ∼0.6). The CFU/ml of the bacilli was determined by plating the cells after serial dilution on Middlebrook 7H10 agar plates and incubating the plates in a bacteriological incubator at 37°C for 2-3 days to get colonies.

*Staphylococcus aureus* ATCC 25923 (Table S1) was cultured in Luria-Bertani (LB) broth and the secondary culture (∼0.6 OD_600_ nm) was used for glycerol stock preparation. *S. aureus* was used for the bioassay to determine the concentration of rifampicin in the culture medium during prolonged exposure of *Msm* to the antibiotic.

*Escherichia coli* JC10289, GM159, JM109 cells (Table S1) were grown in LB broth for the propagation and preparation of plasmids for generation of *Msm arr*-KO and complementation of *arr*-KO with *Msm arr*.

### Generation of *Msm arr* knockout mutant (*Msm arr*-KO)

The *Msm arr* (MSMEG_1221) knockout (*arr*-KO) mutant was generated using the allelic exchange method, as described (37). Experimental details are explained in the Supplementary Materials and Methods.

### Generation of *arr*-KO strain complemented with genome-integrated *Msm arr*

pMV306 (71; Table S1), a genome integrant vector was used for the complementation of the *arr*-KO strain with *Msm arr* under its own promoter. Experimental details are explained in the Supplementary Materials and Methods.

### Growth characteristics of the *Msm* WT and *arr-*KO strains

The *Msm* WT and *arr*-KO strains were inoculated (1%) in rifampicin-free Middlebrook 7H9 broth. The OD_600 nm_ were taken once every 3 hrs using UV-1800 UV-VIS Spectrophotometer (Shimadzu). The growth curves obtained were used for the calculation of mass doubling time of the respective strain. The mass doubling time was calculated from the log phase of the growth curve. Two points on the log phase were selected and the following formula was applied (72):

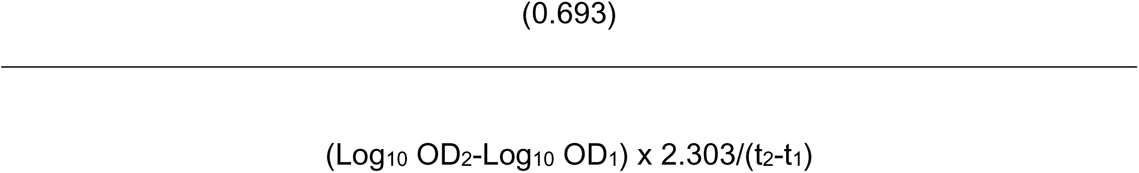

For two OD values chosen on the line, OD1 and OD2, the corresponding time points are t1 and t2, respectively.

Detection and quantitation of hydroxyl radical and superoxide in the MLP cells of the WT, *arr*-KO, *arr*-KO/pMV306-VC, and *arr*-KO/pMV306-*Msm*-*arr* strains

To detect hydroxyl radical and superoxide formation in the MLP cells of WT, *arr*-KO, *arr*-KO/pMV306-VC, and *arr*-KO/pMV306-*Msm*-*arr* strains, the hydroxyl radical specific fluorescence reporter dye, 3’-(p-hydroxyphenyl) fluorescein (HPF, Cayman chemical) (42), and superoxide specific fluorescence reporter dye, CellROX Green (Invitrogen) (46, 47), respectively, was used. Experimental details are explained in the Supplementary Materials and Methods.

### Determination of MBC of rifampicin and moxifloxacin for *Msm arr*-KO strain

*Msm arr*-KO MLP cells were incubated with two-fold increasing concentrations of rifampicin (1.5625 µg/ml to 100 µg/ml) and moxifloxacin (0.125 µg/ml to 8 µg/ml) for a period of 12 hrs, under shaking condition at 170 rpm in a bacteriological incubator shaker at 37°C, as described (50). Subsequently, the cells were subjected to mild sonication of 3 pulses of one-second duration each with one-second interval at 16% amplitude using microprobe in Vibra Cell (Sonics and Materials Inc., USA). The sonicated samples were serially diluted and plated on Middlebrook 7H10 agar plates before and after the antibiotic exposure. For plating of the dilutions below 10^-2^, the aliquots were washed once with Middlebrook 7H9 broth to avoid any antibiotic carryover. The 1x MBC was considered as the lowest concentration of the antibiotic that leads to 2-log_10_ units of reduction in the CFU/ml during the period of incubation, as defined (73).

### Rifampicin bioassay

*Staphylococcus aureus* ATCC 25923 (Table S1) was used for the determination of the concentration of rifampicin (51), with minor modifications as described (36). Fifty µl of rifampicin-exposed culture supernatant from various time points were added into the wells in the *S. aureus* ATCC 25923 embedded in LB agar and incubated in upright position overnight (13 hrs) at 37°C, in bacteriological incubator. The zone of inhibition diameter was measured using Vernier calipers. The standard graph was prepared using known concentrations (0.1 µg/ml to 1 µg/ml, with 0.1 µg/ml interval in concentration) of rifampicin, and used for determination of the rifampicin concentration in the medium of experimental samples. For higher concentrations, solutions were diluted to obtain a concentration in the sensitive range of the assay.

### Susceptibility profile of rifampicin-exposed WT and *arr*-KO strains

Rifampicin (50 mg/ml) (MP Biomedicals) solution was prepared from the dry powder sample in dimethyl sulfoxide (DMSO) and filter-sterilised using Millex-GV PVDF syringe filtration unit of 0.22 µm. The filter-sterilised rifampicin solution was added into *Msm* WT and *arr*-KO MLP culture in biological triplicates to get a final concentration 75 µg/ml and 4 µg/ml (∼2x MBC for respective strains), respectively. Rifampicin susceptibility and the selection of resistant mutants for *Msm* WT and *arr*-KO were obtained by plating on Middlebrook 7H10 in the absence and presence of 125 µg/ml and 6 µg/ml of rifampicin (∼3x MBC for the respective strains), respectively. Experimental details are given under Supplementary Materials and Methods.

### Determination of mutations in the rifampicin-resisters of the WT and *arr*-KO strains

Rifampicin-resistant colonies were isolated from the ∼3x MBC (125 μg/ml) rifampicin plates of the WT and the *arr*-KO strains and inoculated into the respective 3x MBC rifampicin-containing Middlebrook 7H9 broth. These cultures were grown under shaking at 170 rpm, at 37°C, till MLP. The genomic DNA from the rifampicin-resisters of the WT and *arr*-KO strains were isolated using phenol:chloroform extraction method, as described (36), with minor changes in the cell lysis procedure as described in detail in the Supplementary Materials and Methods. The rifampicin resistance determining region (RRDR) of the *rpoB* gene was amplified from the genomic DNA samples of the rifampicin-resisters, control samples (of WT and *arr*-KO cells unexposed to rifampicin) using RRDR-specific primer pairs, Msm-rpoB-RRDR-f and Msm-rpoB-RRDR-r (Table S2) and Phusion DNA polymerase (Thermo Fisher Scientific).The PCR products from the RRDR locus was gel-purified using GeneJet Gel Extraction Kit (Thermo Fischer Scientific) and sequenced on both the strands. The sequencing reactions were performed by M/s. Chromous Biotech, Bangalore, India, or Biokart, Bangalore, India, using both forward and reverse primers of RRDR region. Multiple sequence alignment of the forward and reverse sequencing reactions of the resisters were performed against the forward and reverse sequences of the respective control sample using Clustal Omega (https://www.ebi.ac.uk/Tools/msa/clustalo/) to identify the mutations. Nucleotide changes, which appeared on both the strands of the *rpoB*, were considered authentic.

### Determination of rifampicin resister generation frequency of the MLP and persister cells of the WT and *arr*-KO strains

The resister generation frequency of the MLP cells of WT and *arr*-KO strains to rifampicin was determined by plating the entire 50 ml of the respective MLP culture on Middlebrook 7H10 agar plate containing the respective ∼3x MBC rifampicin. One ml aliquot of the culture was sonicated for 3 pulses of one second each with one-second interval at 16% amplitude using micro-probe of Vibra Cell (Sonics and Materials Inc., USA), serially diluted and plated to determine CFU/ml. The resister generation frequency was calculated using the following formula: Number of antibiotic resisters divided by the total number of the cells from 50 ml culture. Similarly, the resister generation frequency was determined for 4x and 5x MBC of rifampicin also, for the WT and *arr*-KO strains, respectively.

For the resister generation frequency of rifampicin persister cells of the WT and *arr*-KO strains, the number of resisters was divided by the total number of cells at each time point of the respective persistence phase time points, from 24 hr to 48 hr for the WT strain and from 24 hr to 42 hr for the *arr*-KO strain. The average of the resister generation frequencies for all the respective time points of the persistence phase was taken as the resister generation frequency of the persistence phase cells of the respective strain.

### EPR spectrometric analysis for the detection and quantitation of hydroxyl radical

The MLP and rifampicin-exposed persistence phase cells of the WT and *arr*-KO strains, in the presence and absence of thiourea (TU; 150 mM of TU was added from 0 hr itself, 74), were harvested, snap-frozen in liquid nitrogen, and lysed using Teflon pestle. The powdered cell lysate was resuspended in 200 µl of 100 mM sodium acetate (pH 5.2) and centrifuged at 12000 x g for 5 min at room temperature. The supernatant was then mixed with 5,5-dimethyl-1-pyrroline N-oxide (DMPO) to obtain a final concentration of 0.1 M, as described (75). Reading were taken for DMPO-OH adducts exactly 2 min after the addition of DMPO. Samples were loaded into an aqueous flat cell (ES-LC12) and analysed in JEOL JES-X3 ESR spectrometer (FA 200) using the following parameters: X-band, frequency-9428.401 (MHz), Power-4.00000 (mW), Field center-337.275(mT), and Sweep time-2.0 (min). EPR signals were obtained at g ≈ 2. Data were processed using Wizard of Baseline and Peaks in Origin® software.

### Flow cytometry analysis for the detection and quantitation of hydroxyl radical

The hydroxyl radical specific fluorescence reporter dye, 3’-(p-hydroxyphenyl) fluorescein (HPF, Life Technologies) (42), was used to detect hydroxyl radical formation during rifampicin exposure of *Msm* cells. Cells from the different stages of rifampicin exposure of the WT and *arr*-KO strains were collected, and used for staining. The staining and flow cytometry ananlysis was carried out as mentioned in the Supplementary Materials and Methods for hydroxyl radical detection from MLP cells.

## Acknowledgements

PA dedicates this work as a tribute to Prof. T. Ramakrishnan (late), who led the pioneering and foundation-laying work on the biochemistry and molecular biology of *Mycobacterium tuberculosis* at Indian Institute of Science, Bangalore. Authors acknowledge DBT-supported FACS facility, Biological Sciences Division, Indian Institute of Science, and are thankful to Dr. Uttara Chakraborty, Mr. Vasista Adiga, Ms. Urvashi Chauhan, and Ms. Leepika Baid for technical advice and help in flow cytometry measurements. IIT Madras, SAIF, is acknowledged for the EPR and authors thank Mr. N. Sivaramakrishnan and Mr. N. Chandrasekaran of IIT-SAIF for technical help in the EPR measurements.

## Funding

DBT-IISc Partnership Programme for funding and DST-FIST, UGC Centre for Advanced Study, ICMR Centre for Advanced Study in Molecular Medical Microbiology and IISc for the infrastructure support. SS, RRN, AP were CSIR-SRF, and JS was DBT-SRF.

## Conflicts of interest

statement None declared.

## Ethical approval

Not required.

## Author Contributions

PA, SS conceived and designed expts; SS, AP, RRN performed expts; PA, SS, AP, RRN analysed data; PA contributed reagents, materials, and analysis tools; PA, SS wrote the manuscript; all the authors have read the manuscript.

